# A high aneuploidy supply rate underlies parallel evolution of fluconazole resistance in *Saccharomyces cerevisiae*

**DOI:** 10.64898/2026.07.12.738117

**Authors:** Saaz Sakrikar, Akriti Agarwal, Ina Suresh, David Gresham

## Abstract

Copy number variants (CNVs) of *ERG11* play a major role in acquisition of resistance to the commonly used azole family of antifungal drugs in many pathogenic yeast species. However, the conditions in which these CNVs arise, and the factors that underlie their emergence and selection in evolving populations, remain poorly understood. Here, we studied the dynamics of *de novo* CNVs at the *ERG11* locus in *Saccharomyces cerevisiae* to determine the effect of different drug concentrations and temperatures on CNV formation and selection. We found that *ERG11* CNVs emerge reproducibly at fluconazole concentrations around the IC_50_. With increases in temperature, CNVs emerged more rapidly and had a higher tendency towards fixation. Evolved *ERG11* CNV strains provided a significant growth advantage across a narrow range of fluconazole concentrations near the IC_50_ of the wildtype strain. Whole genome sequencing of 35 independent evolved lineages revealed that all *ERG11* CNVs are aneuploidies of chromosome VIII. Relocation of *ERG11* to chromosome XI also results exclusively in selection for chromosome XI aneuploids. We show that fluconazole does not increase the frequency at which aneuploids are generated. Therefore, we conclude that a high spontaneous rate of aneuploidy formation underlies recurrent acquisition of resistance to fluconazole in a temperature and drug concentration dependent manner.

**Author Summary:** Copy number variations (CNV), defined as duplications or deletions of genomic regions, are pervasive throughout all forms of life. Although CNVs can provide a route toward rapid adaptation, they can also be associated with high fitness costs. In pathogenic yeast species, CNVs with diverse genetic structures are known to play a role in antifungal drug resistance. However, the conditions that favor CNV formation and selection are unclear. We applied a fluorescent CNV reporter system to track newly formed CNVs and found that they arise reproducibly at a narrow range of drug concentrations, and that increases in temperature generally lead to more rapid emergence of CNVs. DNA sequencing revealed that the evolved CNVs contained duplications of the entire chromosome, rather than just the region surrounding the gene under selection. Thus, aneuploidy - gain or loss of an entire chromosome - is a mechanism for rapid evolution of drug resistance, potentially without the fitness costs thought to be associated with this mechanism.

## Introduction

Copy number variants (CNVs) are a class of mutation that drive adaptation in diverse species [1,2]. CNVs include amplification or deletion of DNA sequences of various sizes, from a small genomic segment to an entire chromosome, known as aneuploidy [2,3]. Across yeast species, such genomic plasticity has been observed by sequencing wild or clinical isolates [4,5], and a variety of adaptive CNVs have been observed when lab strains are exposed to stresses such as nutrient limitation and antifungal drugs [3,6,7].

CNVs can be the source of both long-term and short-term adaptation. Over the long-term, gene duplication, followed by neo-functionalization of the duplicated copy is a major source of genome evolution [8]. In the short-term, CNVs can lead to an immediate increase in gene expression of one or multiple genes, and hence confer a selective advantage when higher expression of any of these genes is beneficial. However, CNVs also impose a cost of additional DNA and increased expression of additional ‘hitchhiker’ genes within the amplified region that may be deleterious in the selective environment. This has led to the proposal that CNVs, especially aneuploidies, are a rapid but transient adaptation to stress [9–12]. Understanding the dynamics, molecular mechanisms, and long-term fate of CNVs is crucial to our understanding of how organisms and genomes evolve in response to selective pressures.

Accurate detection of *de novo* CNVs in evolving populations, and quantification of their allele frequencies, typically requires isolating and sequencing individual clones. The development of a reporter system for CNVs at a specific locus enables tracking of the emergence and dynamics of CNVs at a single-cell resolution within a mixed population. The CNV reporter consists of a constitutively-expressed fluorescence protein gene located proximal to the gene of interest. Previously, this system has been used to track amplifications of the *GAP1* locus in *S. cerevisiae* in long-term experimental evolution in nitrogen-limiting conditions [6,13] and CNV dynamics in response to antifungal drugs in *Candida albicans [12]*. CNVs detected by an increase in fluorescence can subsequently be confirmed using DNA sequencing to resolve their molecular structures.

Although experimental evolution studies have shown that CNVs reproducibly arise in specific selective conditions, it is unclear how the strength of selection impacts the emergence and selection of CNVs. In the case of experimental evolution in nutrient-limited chemostats, CNVs result in increased expression of nutrient transporter genes enabling CNV-containing cells to out-compete wildtype cells. By contrast, CNVs containing transporter genes are not typically selected when strains are propagated in batch culture conditions even in suboptimal nutritional conditions [14,15].

*ERG11* provides a potential model system to study the determinants of CNV formation and dynamics. *ERG11*, also known as *CYP51* in some species, is an essential gene present across yeast species that encodes the lanosterol 14-α-demethylase enzyme, part of the ergosterol biosynthesis pathway [16,17]. Azole drugs, the most commonly used antifungals, inhibit 14-α-demethylase activity and disrupt the ergosterol pathway, causing an accumulation of toxic intermediates, and reducing cell membrane permeability [18,19]. CNVs of *ERG11,* causing an increase in 14-α-demethylase expression, present a rapid pathway to azole resistance in yeast strains.

*ERG11* CNVs were first observed in treatment-resistant *Candida glabrata* in the late 1990s [4]. They have subsequently been extensively characterized in *Candida albicans* using both clinical samples [10,20] and experimental evolution [7,21,22]. In *Candida albicans*, cells with an isochromosome structure consisting of the left arm of chromosome V, which contains *ERG11,* were found in both clinical samples [20] and after experimental evolution in fluconazole [21], and it was further determined that *ERG11* (along with the transcription factor *TAC1*) was responsible for the enhanced resistance of this strain [23]. *ERG11* amplifications of all types - including aneuploidies, segmental duplications, and local amplifications - have also been found in other *Candida* species, including *C. auris*, and *C. tropicalis* [24–27]. In *Cryptococcus neoformans,* a Basidiomycete species, it was found that aneuploidies contribute to a phenomenon known as hetero-resistance, in which a subpopulation of cells exhibit unstable and reversible resistance to fluconazole [28–30]. Amplification of both *ERG11* and a drug transporter *AFR1* are required for this phenotype [30]. Work in both *C. albicans* and *C. neoformans* has suggested that exposure to fluconazole may stimulate aneuploidy formation, potentially through multimeric cell intermediates that arise due to cell division defects [31–33]. Amplification of *ERG11* is not the only mechanism of fluconazole resistance. Multiple surveys of clinical samples and experimental evolution studies across diverse *Candida* species have not identified *ERG11* amplifications, and have found that other CNVs or SNV-based mechanisms occur more frequently. [34–37]. These contrasting results may reflect differences in experimental conditions as one study in *C. albicans* found that *ERG11* CNVs are repeatedly selected at lower fluconazole concentrations, but were less common at supra-MIC levels [22]. Experimental evolution of *Saccharomyces cerevisiae* in fluconazole revealed SNPs in drug transporters and within the ergosterol biosynthesis pathway, but did not report *ERG11* CNVs [38,39] despite the fact that engineered overexpression of *ERG11* does increase resistance to fluconazole [40].

Thus, although *ERG11* CNVs are found repeatedly in many fungal species, the extent of conservation of this mechanism across the clade is unclear. Furthermore, the specific conditions under which they are selected have not been systematically investigated. Prior experimental evolution studies have been carried out at different temperatures and fluconazole concentrations, both of which are evolutionarily- and clinically-relevant parameters. To address the conservation of *ERG11* amplification as an evolutionary mechanism of adaptation to fluconazole exposure, and to systematically investigate factors that impact its occurrence, we constructed a fluorescent reporter system to track *ERG11* CNVs in the non-pathogenic model species *Saccharomyces cerevisiae*. We carried out experimental evolution for ∼30-105 generations using independent populations of the *ERG11* CNV reporter strain, over a range of fluconazole concentrations and temperatures, allowing us to ascertain CNV emergence and dynamics. The proportion of *de novo ERG11* CNVs were tracked by flow cytometry every passage, corresponding to ∼3-6 generations.

We found that *ERG11* CNVs are selected in multiple independently evolving populations grown in the presence of fluconazole. However, the selective advantage of *ERG11* CNVs is restricted to a narrow range of fluconazole concentrations and temperatures that result in mild or moderate growth inhibition. Surprisingly, we found that all *de novo ERG11* CNVs are aneuploidies, rather than local amplifications. Translocating *ERG11* to the *GAP1* locus on chromosome XI, a locus known to support local genomic amplifications, also resulted exclusively in selection for aneuploidies. By measuring the spontaneous rate of CNV formation and the mechanisms of their formation in the presence and absence of the drug, we show that fluconazole exposure does not result in an increased rate of aneuploidy formation. Therefore, we conclude that the observed highly parallel evolution of *ERG11* amplification to confer fluconazole resistance is attributable to the intrinsically higher rate of aneuploidy compared with other classes of CNVs.

## Results

### *ERG11* CNVs are generated and selected in multiple fluconazole and temperature conditions

We sought to test the evolutionary dynamics of *ERG11* CNVs in response to sustained fluconazole exposure by carrying out experimental evolution using a CNV reporter system. Cells were modified to express the *ERG11* CNV reporter by integration of a constitutively expressed mCitrine gene proximal to *ERG11* (**Fig 1A, S1A Fig**). Experimental evolution of this strain was carried out using serial dilution in batch cultures (**Fig 1B**). Cultures were grown in fluconazole for 2-3 days, then diluted into fresh media containing the same concentration of fluconazole. We repeated these cycles for between 3-6 weeks, corresponding to 10-20 passages or ∼30-105 generations. At every dilution, flow cytometry was performed using a sample of the population to detect *de novo ERG11* CNVs. CNVs were detected through increased mCitrine fluorescence using gating based on 1- and 2-copy controls [41]. Putative *ERG11* CNV-containing lineages from each population were archived at −80℃ for further analysis.

**Fig. 1.**
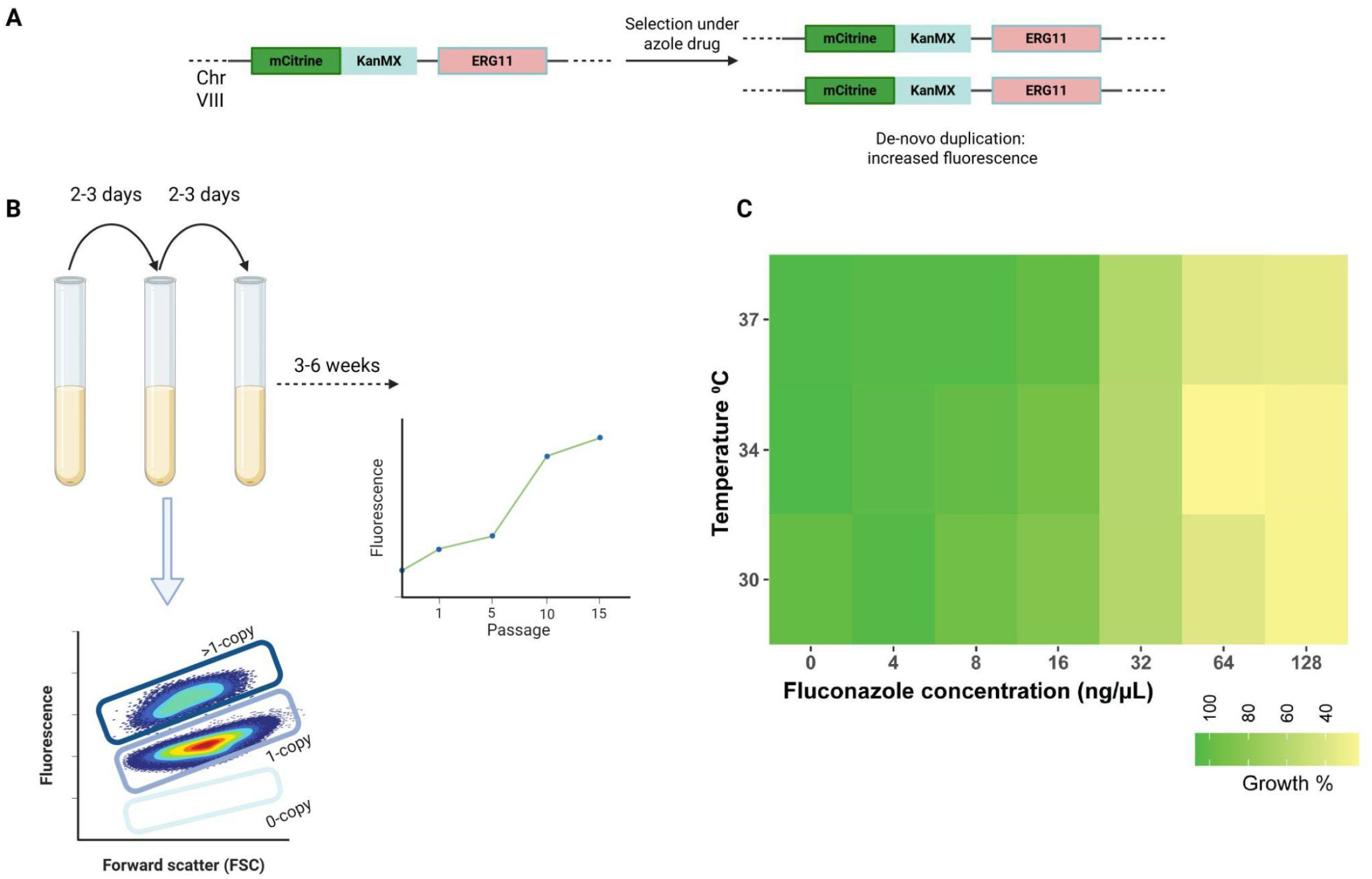
Tracking *ERG11* CNV evolution across a range of fluconazole concentrations. (**A**) The *ERG11* CNV reporter consists of a constitutively expressed mCitrine gene inserted proximal to *ERG11*. (**B**). Cells were grown in a range of fluconazole concentrations (0, 4, 8, 16, 32, 64, 128ng/μL) and diluted between 1:20 and 1:100 every 2-3 days (1 passage) into the same fluconazole concentration. At each dilution, mCitrine fluorescence was measured using flow cytometry. Serial dilutions continued for 20-40 days (9-16 passages) at 3 different temperatures (30℃, 34℃, 37℃). CNVs are observed as an increase in fluorescence over time. (**C**) Fitness landscape of wildtype *S. cerevisiae* over a range of fluconazole concentrations and temperatures, with growth, measured as area under the curve (AUC) relative to 30℃ with no fluconazole.

To determine appropriate selective conditions, we carried out growth assays of *S. cerevisiae* across a range of fluconazole concentrations and temperatures. Drug concentrations were chosen based on values reported in the literature, whereas temperatures were chosen to correspond to temperatures for optimal yeast growth (30℃), human skin (34℃), and human body (37℃). We found that at concentrations less than 16ng/µL, population growth is comparable (94 - 106%) to no-drug control at all three temperatures (**Fig 1C**). By contrast, there is a substantial decrease inf growth at 32ng/µL (56 - 60% of no-drug control). At the higher concentrations of 64 ng/µL and 128 ng/µL, growth was reduced further (21-37%), and inspection of cells under the microscope after 24 hours of growth revealed a substantial amount of cellular debris. We carried out experimental evolution at each combination of temperature and drug concentration to systematically define the selection space for *ERG11* CNVs.

*ERG11* CNVs were detected in environments containing between 8 - 32ng/µL of fluconazole (**Fig 2A** and **S2 Fig**). We observed no CNVs in the absence of fluconazole selection nor at the lowest fluconazole concentration tested (4ng/µL). At 8ng/µL, CNVs visibly emerged around passage 6 at 30℃ and rose to between 20-60% of the population by the end of the run (see 8ng/µL inset in **Fig 2B**). However, no CNVs were seen at this concentration at 37℃.

**Fig. 2.**
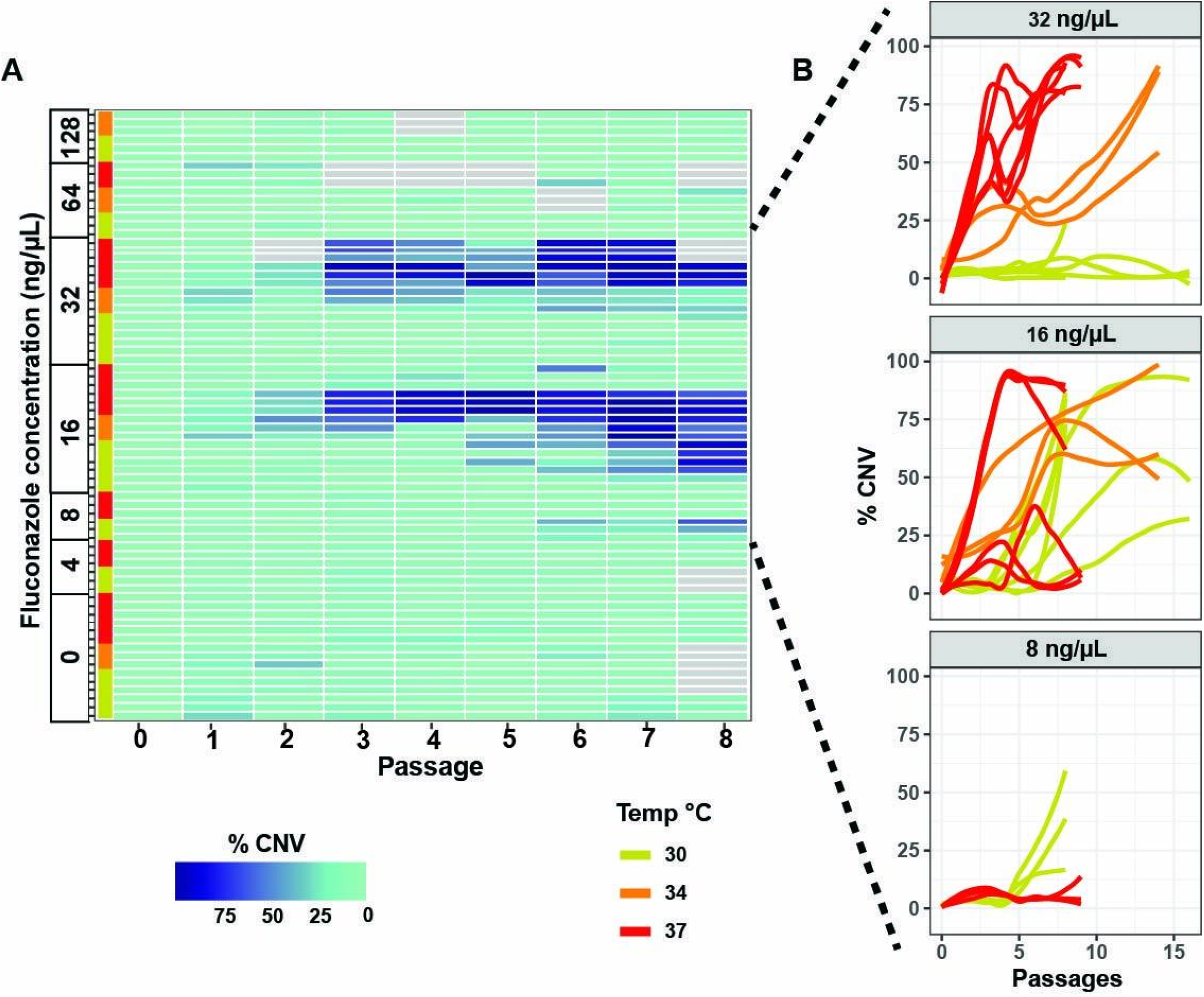
*ERG11* CNVs arise in a temperature- and drug concentration-dependent manner. (**A**) Heatmap, in which every row is an independent population and every column is a passage of serial dilution, of the estimated percentage of *ERG11* CNVs in the population denoted by shade of blue color. Labels and the vertical line (shades of red) denote the fluconazole concentration and temperature at which the experimental evolution was carried out. Data is limited to the initial 8 passages. Gray denotes passages for which data was not obtained. (**B**) Line graphs of *ERG11* CNV dynamics over complete experimental evolution runs at the three fluconazole concentrations in which CNVs were observed. Each line represents an independent population and line color denotes temperature at which populations were evolved.

At 16ng/µL, we observed CNVs at all three temperatures (**Fig 2B**, 16ng/µL inset). Five of 6 independent populations at 30℃ contained CNVs, emerging between the fifth and eighth passages with a final proportion of between 20% and near-fixation (>95%). At 34℃, CNVs were seen in 3/3 independent populations. Interestingly, they emerged much faster (prior to the 2nd passage), and constituted between 50-100% of the population at the end of the run. Similarly, at 37℃, we observed rapid CNV emergence in 4/6 populations, within 2-5 passages. For three of these populations, CNV-containing lineages formed the majority of the population (60-85%) at the end of the run.

Temperature appeared to accelerate the emergence of *ERG11* CNVs. The effect of temperature on CNV dynamics was most clear at 32ng/µL (**Fig 2B**). CNV emergence was rapid at the highest temperature 37℃ first appearing within 3 passages, and 6/6 populations near fixation (80-100%) within 3-8 passages. At 34℃, CNV emergence was observed in all 3 populations between 2-5 passages, and 2 of 3 populations were approaching fixation of the CNV after 14 passages. However, at 30℃, no CNV was seen in any of 6 independent populations. Thus, *ERG11* CNVs emerged only at concentrations between 8 and 32ng/µL, and we observed no CNVs at the higher fluconazole concentrations of 64 and 128ng/µL. Notably, the concentrations at which CNVs arise bookend the IC_50_ value we obtained for the wildtype parental *S. cerevisiae* strain in rich media (**S3 Fig**).

### *ERG11* CNVs confer growth advantage at specific fluconazole concentrations

To define the benefits and costs of *ERG11* CNVs we studied the growth characteristics of the evolved strains. We isolated a total of 35 *ERG11* CNV-containing lineages from 18 independent populations based on increased fluorescence (see **Methods**). From each unique condition of temperature and drug concentration in which CNVs evolved, at least two strains (a total of 21 strains) were randomly chosen for phenotyping.

*ERG11* CNV-containing lineages were grown for 24 hours in 0, 8, 16, 32, and 64ng/µL fluconazole at 30℃. Growth rates were calculated for each strain in each concentration of fluconazole by fitting data to a logistic curve. At each fluconazole concentration, the growth rate for each CNV strain was then normalized to the average growth rate for control strains in the same conditions (**Fig 3A)**.

**Fig. 3.**
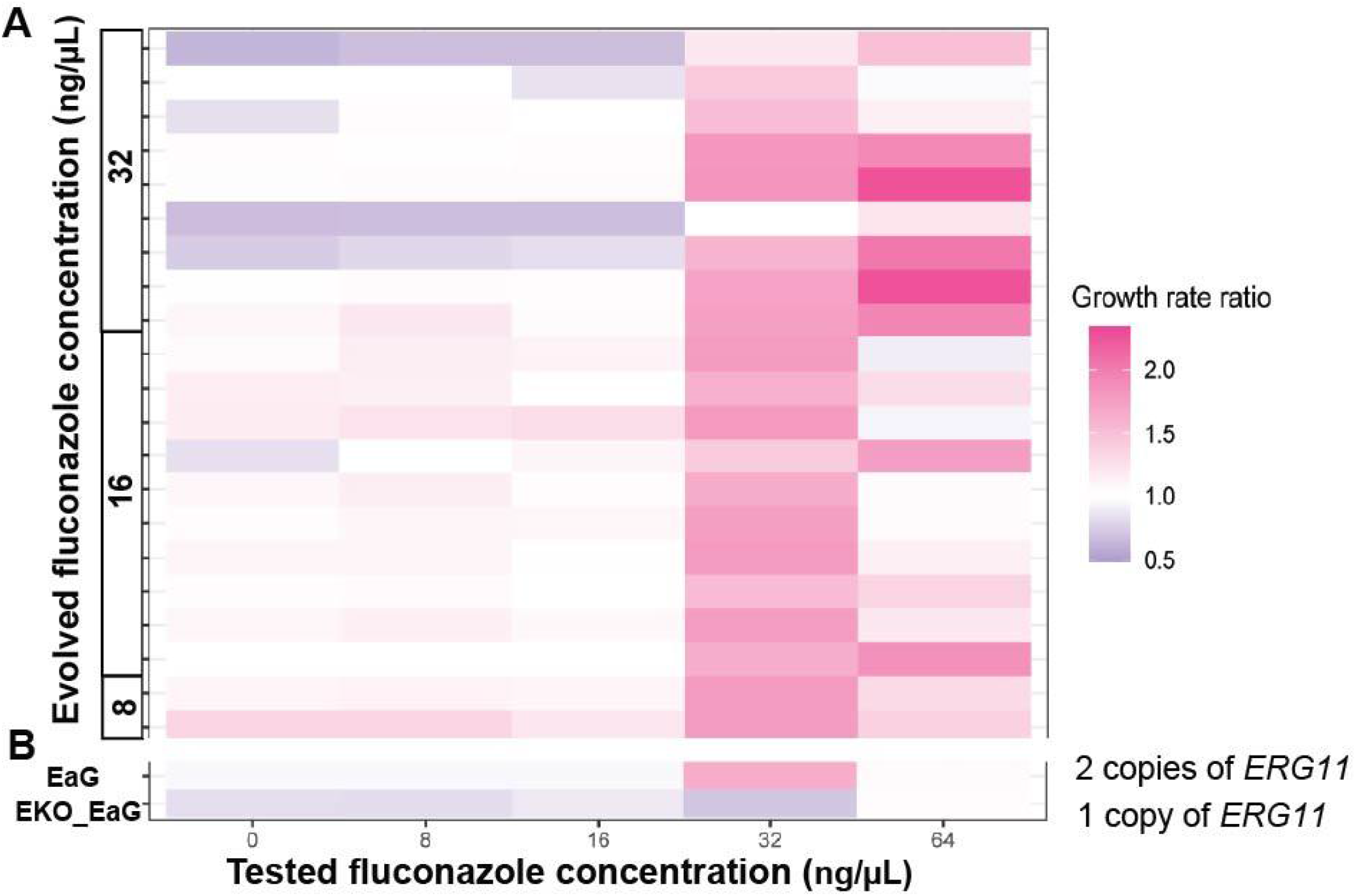
An extra copy of *ERG11* provides a fitness benefit predominantly at 32ng/µL fluconazole. In the heatmap, each row represents a different strain, and each column denotes a different fluconazole concentration. The growth rate ratio of each strain is depicted by the heatmap colors, with respect to the growth rate of control strains at that concentration. (**A**) Evolved *ERG11* CNV strains,sorted by the concentration at which they were evolved, and (**B**) engineered strains containing two (EaG) or one (EKO_EaG) copies of *ERG11*.

First, we assessed whether the presence of *ERG11* CNVs reduces growth in the absence of the selective condition (first column of **Fig 3A)**. Overall, the average growth rate of the CNV strains without fluconazole was not significantly different from controls (*p*=0.11, t-test). However, seven strains did grow significantly slower than the control in the absence of fluconazole (t-test, p< 0.05). All seven were found to have evolved at elevated temperatures, whereas the growth assay was carried out at 30℃. We conclude that most of the evolved CNVs do not adversely impact growth in the absence of the selective pressure of fluconazole.

Next, we assessed growth of *ERG11* CNV lineages in the presence of fluconazole, at the drug concentration at which the evolution was performed. The two CNV strains evolved at 8ng/µL fluconazole grew at 121% of control strain rate at that concentration (*p*<0.008, one-tailed t-test). The ten CNV strains evolved at 16ng/µL of fluconazole grew at 106% of control strain growth rate at that concentration (*p*<0.004, one-tailed t-test). The nine CNV strains evolved at 32ng/µL of fluconazole grew at 149% of control strain growth rate at the corresponding concentration (*p*<3e-10, one-tailed t-test). Looking at strains individually, 10/21 strains demonstrate significantly higher growth rate than the controls at the fluconazole concentration in which they were evolved (t-test, p < 0.05). The remaining strains had higher growth rates than the controls, but the differences were not statistically significant. Although, it is unexpected to find that some strains do not show a significant advantage at the concentration at which they were evolved this may partly be explained by the fact that growth assays were carried out over 24 hours, while strains were evolved with serial dilution every 48-72 hours, and that growth assays were carried out at 30℃, while some CNV strains were evolved at 34 or 37℃.

Notably, we observed the strongest growth advantage at 32ng/μL fluconazole for all evolved strains. Regardless of the fluconazole concentration at which strains had evolved, 19 of the 21 tested strains showed a significant growth advantage at 32ng/μL (t-test, p < 0.05). Across all 21 evolved strains, the average growth rate was 162% of the control strains’ growth rate at this concentration. These results suggest a common selective advantage conferred by *ERG11* CNVs, with lineage variation likely resulting from additional variation acquired during the experimental evolution.

### Evolved *ERG11* CNV strains are chromosome VIII aneuploidies

We sought to understand the molecular basis of evolved *ERG11* CNVs using whole-genome sequencing. (see **Methods**). Using sequence read depth analysis, we found that all *ERG11* CNV lineages contain full-chromosome aneuploidies of chromosome VIII, which contains *ERG11* (**Fig 4** and **S4 Fig**). All lineages contained a duplication of chromosome VIII except a single case from a population evolved at 34℃ and 32ng/µL fluconazole, where a triplication of chromosome VIII was detected, consistent with the CNV reporter fluorescence from that lineage **(S4 Fig)**.

**Fig. 4.**
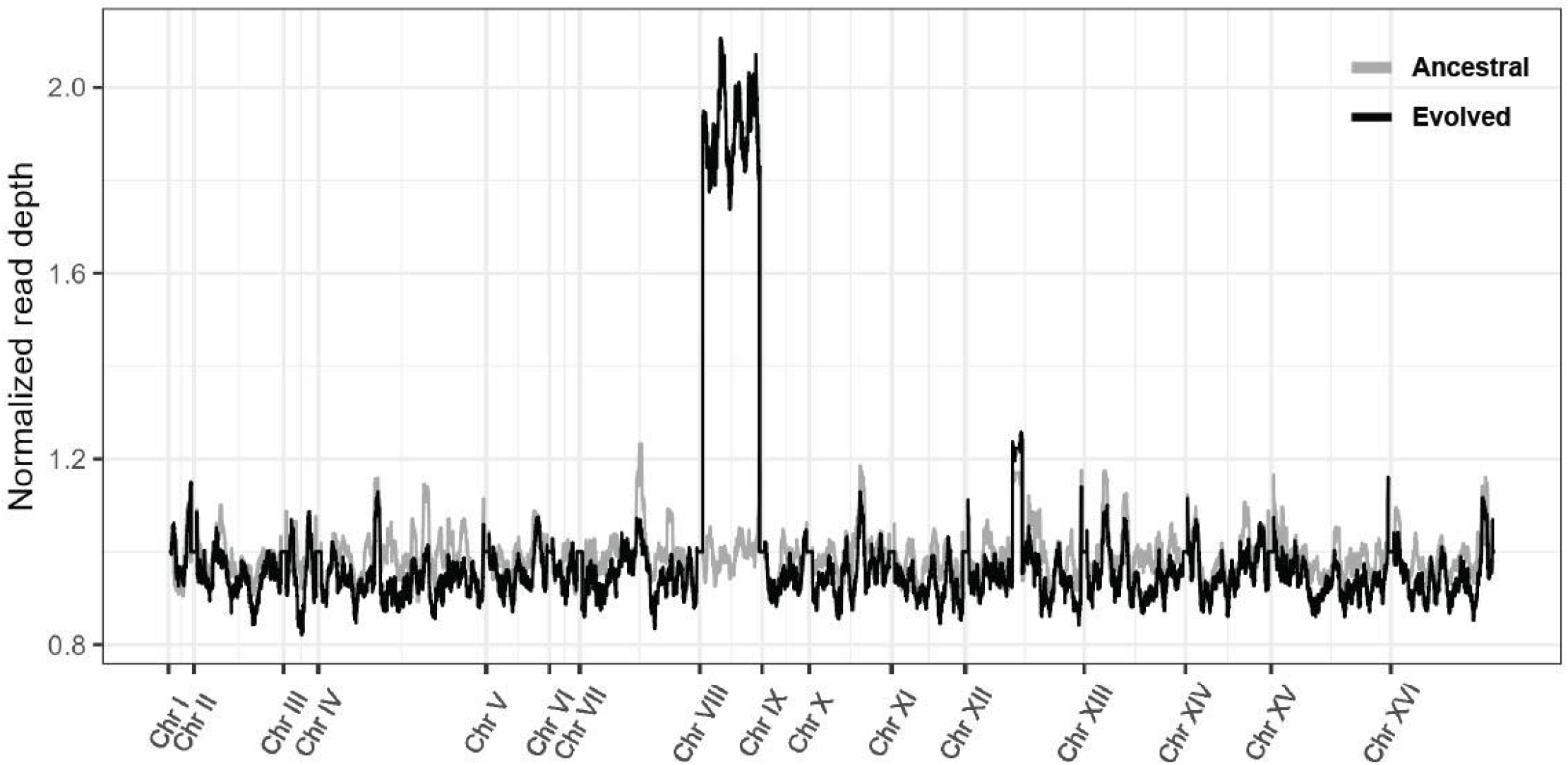
*ERG11* CNV strains are chromosome VIII aneuploidies. Sequence read depth of a representative *ERG11* CNV strain (black) showing twice the read depth on chromosome VIII compared to the read depth of the ancestral strain (grey).

This preponderance of aneuploidies was unexpected, given the diversity of CNVs in *S. cerevisiae* that have been observed in nutrient-limited chemostats, and when *C. albicans* adapts to azole drugs in laboratory and clinical settings [6,22,34,42–44]. Hence, we investigated factors that might explain the parallel evolution of fluconazole resistance through aneuploidy.

### Increased expression of multiple genes throughout the genome increases fitness in fluconazole

To test if an additional copy of *ERG11* explains the growth advantage of selected lineages, we created a strain (EaG) containing an additional copy of *ERG11* on chromosome XI at the *GAP1* locus (**S1B Fig**, see **Methods**). The presence of an extra copy of *ERG11* results in an increased growth rate in fluconazole, most clearly at 32ng/µL of fluconazole (**Fig 3B**). At this concentration, growth of EaG is statistically indistinguishable from evolved CNV strains (pairwise t-test, *p*=0.32). We then deleted the native copy of *ERG11* on chromosome VIII in EaG resulting in a strain, EKO_EaG containing *ERG11* at the *GAP1* locus on chromosome XI only. Deleting the native copy of *ERG11* from EaG abolishes the growth advantage in fluconazole (**Fig 3B**), and, at 32ng/µL, this strain grows significantly slower than evolved CNV strains and the strain EaG (pairwise t-test, *p*<9e-6 and *p*<3e-4 respectively).

Repeated selection for chromosome VIII aneuploidy may be due to genes in addition to *ERG11* on the chromosome VII that increase fitness when amplified. Therefore, we sought to test whether increased expression of additional genes on chromosome VIII increases fitness using the YETI overexpression collection [45]. In addition to *ERG11*, overexpression of additional genes on chromosome VIII, including the lipid regulator *NEM1*, golgi regulator *GGA2*, the mitochondria-associated *TRR2* and *QCR10*, ribosomal Rpl27A, and the RNA Pol subunit *RPC10* result in increased fitness in fluconazole (**Fig 5**). Notably, overexpression of several genes on other chromosomes also increase fitness, including the drug transporter genes *FLR1* and *PDR5*, the stress-responsive transcription factor gene *YAP1*, and *ERG13*, which encodes an additional component of the ergosterol pathway targeted by fluconazole.

**Fig. 5.**
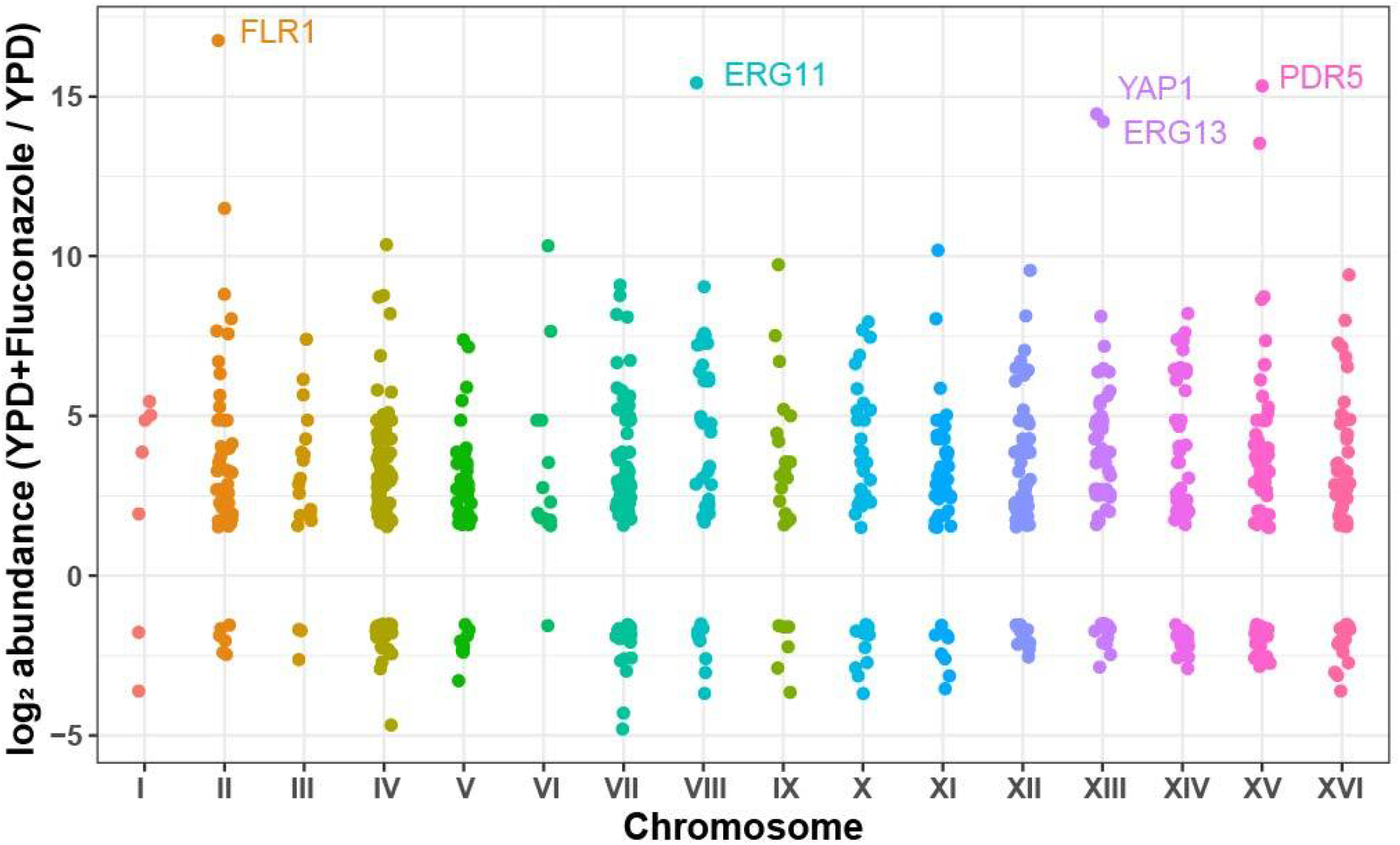
Overexpression of multiple genes increases fitness in fluconazole. Fitness effect of the overexpression of 781 genes that significantly increase fitness during 72 hours exposure to 32ng/µL fluconazole when over-expressed. The relative change in abundance of each strain in fluconazole compared with no fluconazole is a proxy for its fitness effect in the drug condition.

Despite clear evidence for additional genes that are beneficial when overexpressed, we did not find additional CNVs or aneuploidies among *ERG11* CNV lineages (**S4 Fig**).This can be explained by the fact that the CNV reporter used in this study was used to track *ERG11*, present on chromosome VIII, and hence only these lineages were chosen for sequencing. It is also possible that fitness costs associated with overexpression of off-target genes on other chromosomes prevented their selection. Finally, estradiol induction of the YETI collection leads to a gene expression increase much greater than two-fold [45], whereas CNVs typically double the copy number, potentially limiting the informativeness of this assay.

### Recurrent aneuploidy formation is not unique to chromosome VIII

To understand if the preference for aneuploidies is driven by the physical location of *ERG11* on chromosome VIII we tested whether the genomic location of *ERG11* affects the type of CNVs formed. We evolved EKO_EaG, containing *ERG11* and the mCitrine CNV reporter at the *GAP1* locus on chromosome XI at 16ng/µL fluconazole and 30℃. CNVs were detected in 3 of 4 independent evolved populations, with CNV emergence visible after between 4 and 10 passages (**Fig 6A**). These dynamics are not dissimilar to the CNV dynamics for *ERG11* at its native locus when evolved under similar conditions (**Fig 2B**). We isolated four *ERG11* CNV lineages from two of the three evolved populations and sequenced them. We detected chromosome XI aneuploidies in all four lineages (**Fig 6B** and **S5 Fig**). This indicates that the preference for aneuploidies is independent of the genomic location of *ERG11*.

**Fig. 6.**
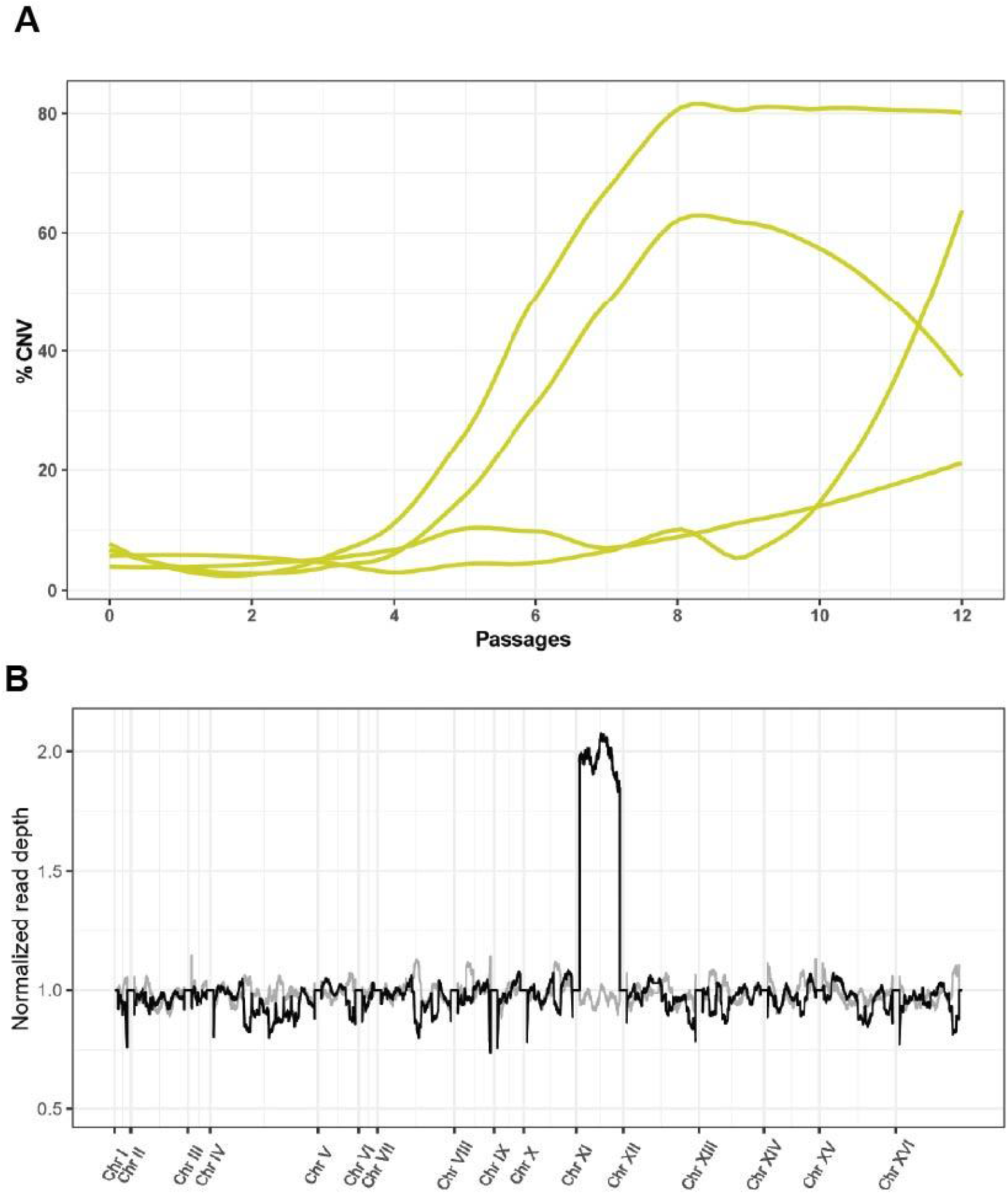
Relocation of *ERG11* to chromosome XI results in chromosome XI aneuploidies during selection for *ERG11* CNVs. (**A**) *ERG11* CNV evolution dynamics for the strain EKO_EaG at 16ng/μL fluconazole and 30℃, and (**B**) normalized read depth for representative CNV strain (black) shown with ancestral strain (grey).

### Fluconazole does not stimulate aneuploidy formation

We sought to test if exposure to fluconazole increases the propensity for aneuploidy formation. To that end, we used a reporter strain, FACu_report, containing one copy each of both *SFA1* and *CUP1* inserted at the *DDI1* locus on chromosome V, with both genes deleted at their native loci [46,47]. *SFA1* encodes formaldehyde dehydrogenase, which degrades toxic formaldehyde, whereas *CUP1* encodes methallothionein, which binds copper ions. Duplication of this locus results in survival on selective media containing formaldehyde and copper sulphate, enabling selection of spontaneous duplications at this locus.

We grew the FACu_report strain in the presence and absence of fluconazole and plated on selective media containing both formaldehyde and copper sulphate, and counted the number of colonies, to obtain an estimated μ, the CNV mutation rate per cell per division (see **Methods**). Using this method, we estimate a spontaneous CNV rate in the absence of fluconazole of 4.3e-5 per cell per generation, compared with 1.4e-5 in 16ng/μL fluconazole and 8e-6 in 32ng/μL fluconazole.

We isolated and sequenced 5 spontaneous *SFA1*-*CUP1* CNVs from media lacking fluconazole and 9 spontaneous *SFA1*-*CUP1* CNVs from media containing fluconazole.. We found that of the 5 strains not exposed to fluconazole, 4 contain chromosome V aneuploidies (**S6A Fig**). In the presence of fluconazole, 6 of the 9 strains contain chromosome V aneuploidies (**S6B Fig**). There is no significant difference in aneuploidy rate between the conditions (Fisher’s exact test, *p*=1.0). In addition, we detected local amplifications of the engineered *SFA1*-*CUP1* locus on chromosome V in 4 fluconazole-exposed and all 5 non-fluonazole-exposed strains (**S6 Fig**). These results indicate that exposure to fluconazole does not stimulate an increased rate of aneuploidy formation in *S. cerevisiae*.

## Discussion

Copy number variation is hypothesized to provide a rapid route for adaptation to stressful conditions [9–11]. Consistent with this hypothesis, amplification of *ERG11* has been seen in a variety of pathogenic fungi in clinical samples and with experimental evolution in the presence of fluconazole [4,12,23,24,26,30]. Here, we demonstrate that the non-pathogenic model species *Saccharomyces cerevisiae* also develops *ERG11* CNVs in response to azole selection. Combined with reports from diverse pathogenic yeast species including *C. albicans*, *C. glabrata*, *C. auris*, and *C. neoformans*, this result confirms that *ERG11* CNVs are a clade-wide mechanism for adaptation to azole drugs.

Within the limited span of our experimental evolutions (estimated at a maximum of ∼105 generations), we found that evolved *ERG11* CNVs were largely stable within the population, and did not get replaced by theoretically less costly adaptations. By contrast, tracking of *C. albicans* patient samples showed CNVs to be a transient or intermediate adaptation [10]. Those were often replaced by strains exhibiting a loss of heterozygosity (LOH), which is not relevant in our haploid species. Furthermore, we note that the majority of our evolved strains did not show a significant growth defect in the absence of selection, suggesting possible long-term stability.

We found that CNVs arose at ∼IC_50_ and sub-IC_50_ concentrations, matching observations from *C. albicans* [22]. As expected, we did not see a notable proportion of *ERG11* CNV cells in the absence of fluconazole selection. We hypothesize that the selective pressure at the lowest drug concentrations was insufficient to see significant amounts of CNV cells in the populations. At higher drug concentrations, even though data from engineered strains suggests an extra copy of *ERG11* can provide a fitness benefit, CNVs were not found. We note that research on *C. albicans* has suggested that exposure to supra-MIC doses of fluconazole tends to select for a tolerant, rather than resistant, phenotype, and that *ERG11* CNVs are not involved in the emergence of tolerance [22]. Hence, it appears that *ERG11* CNVs arise within a limited range of fluconazole concentrations that inhibit growth, and they are not selected below or above this range.

This is the first systemic look at how temperature affects CNV formation. There is a marked effect of temperature on CNV emergence and dynamics, with CNVs emerging within just 2–5 passages at higher temperatures, and populations exhibiting an increased propensity of fixation. It has been reported in *Cryptococcus deneoformans* that even small increases in temperature promote genomic instability, with increased rates of transposon activity [48]. In *S. cerevisiae*, acute stress, including heat shock at much higher temperatures were found to dramatically increase the rate of chromosomal rearrangements, including the gain and loss of entire chromosomes [49,50]. Work in *Arabidopsis thaliana* showed that chronic exposure to milder heat stress results increased crossover rates and impaired chromosome segregation during meiosis [51,52]. Hence, we suggest that the genomic instability caused by higher temperatures allows more rapid and widespread formation of CNVs. This has potential clinical implications, since CNVs were found most readily at human body temperatures (37℃).

We found a significant growth advantage for all *ERG11* CNV strains at the particular concentration of 32ng/μL, regardless of the concentration at which they were evolved. Furthermore, we found that inserting a second copy of *ERG11* in an otherwise native background also provides a similar growth advantage, suggesting that the fitness benefit of the evolved CNVs is due to the extra copy of *ERG11*. It is therefore interesting that all CNV strains were found to be whole-chromosome aneuploids of chromosome VIII rather than local amplifications.

We considered several possible explanations for this observation. First, we observed that genes other than *ERG11* on chromosome VIII provide a fitness advantage when overexpressed in fluconazole (**Fig 5**). It is thus possible that chromosome VIII aneuploidies are fitter than local CNVs because of the combined effect of *ERG11* duplication coupled with the duplication of additional genes on chromosome VIII. Second, it is possible that aneuploidies arise spontaneously at a higher frequency than local CNVs, and are thus more readily selected when the selective pressure of fluconazole is introduced. Third, previous work in *C. albicans* and *C. neoformans* has suggested that the introduction of fluconazole itself stimulates aneuploidy formation, possibly by causing the formation of multimeric cells [30–33]. Stress-stimulated amplification leading to adaptation, via a different mechanism, has also been reported at the highly variable *CUP1* locus of *S. cerevisiae* [53].

We found that adding an extra copy of *ERG11* at a non-native locus provided a growth advantage similar to that seen for the evolved CNV strains, and that removing the native copy abolished this advantage. This indicates that the copy number of *ERG11* alone is sufficient to explain the observed fitness increase in fluconazole. Experimental evolution of a strain containing *ERG11* at the *GAP1* locus on chromosome XI resulted in chromosome XI aneuploidies. Therefore, we conclude that a preference for aneuploidies is not driven by additional beneficial genes on chromosome VIII.

Using an assay for spontaneous CNV formation rates, we did not observe a higher rate of CNVs, nor a bias towards aneuploidies upon acute exposure to fluconazole. Furthermore, in some evolving populations, *ERG11* CNVs were detected in the population only after 8 passages, corresponding to roughly 50 generations of growth, which would be unlikely in a scenario where aneuploidies were induced or stimulated by acute exposure to fluconazole. Finally, a potential mechanism for fluconazole-induced aneuploidy formation reported in other yeast species - multimeric cells following acute fluconazole exposure - was previously tested and not observed in *S. cerevisiae* [32]. Hence, there is strong evidence against the third explanation: that fluconazole directly stimulates aneuploidy formation.

Therefore, we conclude that the striking parallelism in the mechanisms of *ERG11* CNV formation is due to the high supply rate of aneuploids. Estimates of mutation rates in *S. cerevisiae*, using mutation accumulation, found a ten-fold higher prevalence of whole chromosome aneuploidies over “large CNVs” [54]. Since mutation accumulation experiments are not affected by selection except in the case of very strongly deleterious mutations, this result would suggest an intrinsically high supply rate of aneuploidies compared to local amplifications.

By contrast with these results, detailed analyses of *GAP1* CNVs evolved in response to nitrogen limitation [6] have identified few aneuploidies and primarily local amplifications. We hypothesize several mutually reinforcing reasons for this difference. Most chromosome VIII aneuploidies did not have a significant growth defect in the absence of selection, potentially reducing selection pressure that might have favored local amplifications. Similarly, a CNV stability assay found that the fitness cost of aneuploid strains depends on the identity of the duplicated chromosome [55,56]. Another important difference compared to the *GAP1* studies is the number of generations: in this study, evolution was performed over ∼90 generations whereas for prior chemostat studies of *GAP1* amplification selection was maintained for 150-250 generations. Finally, chemostat studies of *GAP1* amplification were performed with a population size of ∼1e8, whereas serial dilution meant that the effective population size in batch cultures, estimated by the method of harmonic means [57], was ∼7e5. The combinations of these factors likely explains the contrasting results between these two evolutionary scenarios.

We note that there are previous studies of azole-resistant CNVs in other yeast species which have produced largely whole-chromosome or segmental aneuploidies [10,12,20,22,30], and others which have produced local amplifications [27,34,58], and that further work could establish the reason for these apparent preferences. In conclusion, our work affirms the potential of aneuploidies to provide a pathway for rapid adaptation to stress, resulting in highly parallel evolutionary outcomes, and shows that growth temperature and strength of selection are key determinants of the dynamics of this adaptative mechanism.

## Methods

### Strains, media and growth conditions

Strains, plasmids, and primers used in this study are listed in **S1 Table**. The *ERG11* CNV reporter strain E_report was created by inserting the mCitrine-KanMX cassette from plasmid DGP71. The cassette consists of mCitrine constitutively expressed under a *ACT1* promoter, coupled with a Kan/G418 resistance cassette (*ACT1pr::mCitrine::ADH1term TEFpr::KanMX::TEFterm*). This cassette was inserted at locus 122500 on chromosome VIII, roughly 1kb upstream of the native *ERG11* locus on chromosome VIII of *S. cerevisiae* FY4 (MATa). The *ERG11* insert strain EaG (*GAP1*Δ::*mCitrine-KanMX-ERG11*) was created from the plasmid DGP481 using a pUC19 backbone. DGP481 was created by adding 500bp homology of the flanking regions of *GAP1* to each end of the *mCitrine-KanMX-ERG11* cassette. This plasmid was transformed into FY4 (MAT a). Finally, the *ERG11* KO strain (EKO_EaG) was created from the insert strain EaG using the HygB cassette of plasmid DGP363, amplified by primers which added 45bp homology to flanking regions of *ERG11.* This cassette was then transformed into EaG. All transformations into yeast were carried out using a modified lithium acetate protocol [59]. Created strains were verified using antibiotic selection, fluorescence, PCR, Sanger sequencing around the modified region, and whole genome sequencing. Fluorescence control strains used here (1_ctrl, 2_ctrl) have been described previously [6].

Cells were grown in YPD media (1% Bacto yeast extract, 2% Bacto peptone, 2% Glucose) for routine culturing, as well as experimental evolution and growth assays. 50mg/mL stocks of fluconazole in DMSO were made and diluted as appropriate for various experiments. In conditions of no fluconazole, indicated throughout the paper as 0ng/µL, DMSO was added to the medium to a final concentration of 0.2% v/v. For the formaldehyde-copper sulphate CNV assay, strain JAY538 (also known as FACu_report) was obtained as a gift from the lab of Juan Lucas Argueso. Non-selective plates were made of SC-Trp dropout media (0.17% Yeast Nitrogen Base, 0.14% Trp dropout mix, 0.5% ammonium sulphate), while selective plates additionally contained 1.8mM formaldehyde and 150μM copper sulphate.

### Experimental evolution by serial dilution

The *ERG11* CNV reporter strain and fluorescence control strain cells were plated onto YPD and multiple independent single colonies were picked and grown in liquid YPD. Overnight cultures were diluted 1:100 into 96-well plates containing YPD supplemented with fluconazole and grown for 48-72 hours, followed by appropriate dilution (between 1:10 and 1:100) into the same fluconazole concentration. At every passage (i.e., every 48-72 hours), aliquots from each well were analyzed with flow cytometry.

These cycles were repeated over 7-16 passages or 20-40 days, corresponding to ∼40-105 generations. For each run, 2-3 independent replicates of the *ERG11* CNV reporter strain and 1-2 replicates each of the 1- and 2-copy control strains were used. The fluconazole concentrations included 0, 4, 8, 16, 32, 64, and 128ng/µL, and the runs were carried out at 30℃, 34℃, and 37℃.

### Flow cytometry

Flow cytometry was carried out using the Cytek Aurora (Cytek Biosciences). The percentage of cells containing CNVs in each sample was estimated by manually gating mCitrine fluorescence, based on the fluorescence levels of the 1- and 2-copy controls, in the SpectroFlo software (Cytek Biosciences). The gate was drawn based on signal from the laser channel B2 (488nm) and the size of the cell measured as forward scatter FSC [41]. The percentage of cells having fluorescence greater than the 1-copy gate were taken as CNV-containing cells. At certain timepoints, the 1- and 2-copy control fluorescence levels showed substantial overlap, making the gating ineffective. In these cases, the CNV percentage was estimated by normalizing the median fluorescence of the CNV reporter strain to the median fluorescence of the control strains. In later runs, the resolution of gating was improved by diluting an aliquot of the evolving cells (1:10 dilution) in YPD, and incubating at 30℃ for ∼6 hours before carrying out flow cytometry. Results from each timepoint are noted in **S2 Table**.

### Isolation of *ERG11* CNV strains

Populations of cells containing CNVs were saved as glycerol stocks, and plated onto YPD plates. Individual colonies were grown overnight in YPD and flow cytometry was performed, to determine the mCitrine copy number. In some cases, YPD plates and liquid media were supplemented with 32ng/μL fluconazole to allow the growth of some otherwise slow-growing populations. In some cases involving mixed populations with a substantial fraction of non-CNV containing cells, or slow growth on YPD plates, a saved population was FACS sorted to isolate a subpopulation of CNV-containing cells. CNV-containing clones were isolated from these sorted populations by plating, growing a single colony overnight in YPD, and verifying increased mCitrine fluorescence.

Colonies exhibiting CNVs (>1-copy of mCitrine) were saved as glycerol stocks, and were subject to phenotyping by growth assays and whole-genome sequencing (see below). A total of 35 CNV-containing strains were isolated, including at least one from each population that showed a CNV. A list of these strains indicating the conditions they were evolved under, is in **S1 Table**.

### Phenotyping CNV strains with growth assays

CNV strains validated based on fluorescence were subject to phenotyping by growth assays in varying concentrations of fluconazole. Strains were grown overnight in YPD (supplemented in some cases with fluconazole), then diluted to an optical density (OD600) of 0.01, and then grown for 24 hours in a 96-well plate in a range of fluconazole concentrations. The plate was placed at 30℃ in the plate reader Tecan Spark (Tecan Trading AG), with OD at 600nm readings recorded every 5 minutes. The resulting growth data was fitted to a logistic growth curve using the R program growthcurver [60], from which the maximal growth rate (*μ*) was obtained. Between 2-7 replicate growth assays were performed for every strain including the control strains. Growth data shown is the average growth rate of the given CNV strain at a given drug concentration (averaged over all replicates), divided by the average growth rate of the control strain at that concentration (averaged over all control replicates). Control strains included multiple replicates each of the *ERG11* CNV reporter strain, 1_ctrl, 2_ctrl, and 0_ctrl.

### Whole-genome sequencing of CNV strains

CNV strains validated by fluorescence were subject to whole-genome sequencing. DNA was extracted using a modified Hoffman-Winston protocol [61] and library preparation was done using a Nextera kit and Illumina adapters. Sequencing was carried out at the NYU GenCore using the Illumina NextSeq 500 platform PE150 (2×150 300 Cycle v2.5) or Illumina NovaSeq 6000 SP PE150 (2×150 300 Cycle v1.5) or Element Bioscience Aviti (2×150).

For alignment of sequencing results, the *S. cerevisiae* R64 reference genome was modified to include the mCitrine-KanMX cassette using Reform (https://gencore.bio.nyu.edu/reform/). The FASTQ files were aligned to the modified genome using bwa [62], and sorted bam files were generated using samtools 1.14 [63]. The sorted BAM files were used to generate per-base depth files using the bedtools genomecov command, and this data was used to plot read depth across the genome.

### YETI overexpression enrichment analysis

The barcoded Yeast Estradiol strain with Titratable Induction (YETI) collection [45], which contains 4963 genes under an estradiol-inducible promoter, was grown in three conditions: 1) YPD, 2) YPD + estradiol, and 3) YPD + estradiol + 16ng/μL fluconazole. Cells were grown for 72hrs and barcode sequencing (Barseq) was performed on cell aliquots at regular timepoints, using a two-step PCR amplification process [64]. Bar-seq results were analysed using DESeq to contrast the barcode abundance in the YPD+Estradiol+Fluconazole versus YPD+Estradiol conditions. Thresholds of padj <0.05, baseMean >50, and |log2FoldChange| > 1.5 were used to determine genes whose overexpression significantly affects fitness in fluconazole.

### Formaldehyde-copper sulphate resistance assay

Single colonies of strain FACu_report were grown overnight (∼24 hours) in YPD supplemented with fluconazole concentrations of 0, 16, and 32ng/μL. Cell counts from overnight cultures were estimated using a hemocytometer, and after appropriate dilutions, 1e5 cells were plated on selective plates (SC-Trp + formaldehyde + copper sulphate), while, for some replicates, 1e2 cells were plates on permissive plates (SC-Trp). These plates were then incubated at 30℃. Counts of colonies on each plate were taken after 48 hours for permissive plates and 72 hours for selective plates. Colony counts on permissive plates were used to establish the accuracy of the hemocytometer-based cell counting. All colony counts are shown in **S3 Table**. 1-2 colonies were randomly chosen from each selective plate and streaked onto permissive medium, and then replica-plated back onto selective medium to establish the stability of resistance to fluconazole/copper sulphate. These stably resistant colonies were then subject to gDNA extraction and sequencing as described above.

### Estimating mutation rates from FACu colony counts

The probability of mutation per cell per generation (μ) was calculated from the estimated number of mutations per culture (m), obtained from the median number of observed mutants (r) in the fluctuation assay, as described [65]. m was obtained from r by solving for the equation *r*/*m* − *ln*(*m*) = 1. 24 originally derived by Lea and Coulson [66]. μ was obtained from m by dividing by the estimated population size at the time of plating (1e5 cells).

## Data availability

Sequencing data is available in NCBI Sequence Read Archive accession PRJNA1419567. Code for data analysis and figures are available at https://github.com/GreshamLab/Saaz_ScERG11CNV

## Acknowledgements

We thank the NYU Center for Genomics and System Biology Genomics Core, especially Carolina Thornton, for their assistance and resources. We thank Juan Lucas Argueso and Ruthie Watson for the generous gift of the strain JAY538 as well as advice regarding its use. We thank Camile Bedard of the Landry lab for advice regarding IC_50_ measurements. We thank current and former Gresham Lab members, in particular Chris Jackson, Pieter Spealman, Titir De, and Julie Chuong, for technical assistance and advice.

## Supplementary figures

**S1 Fig.**
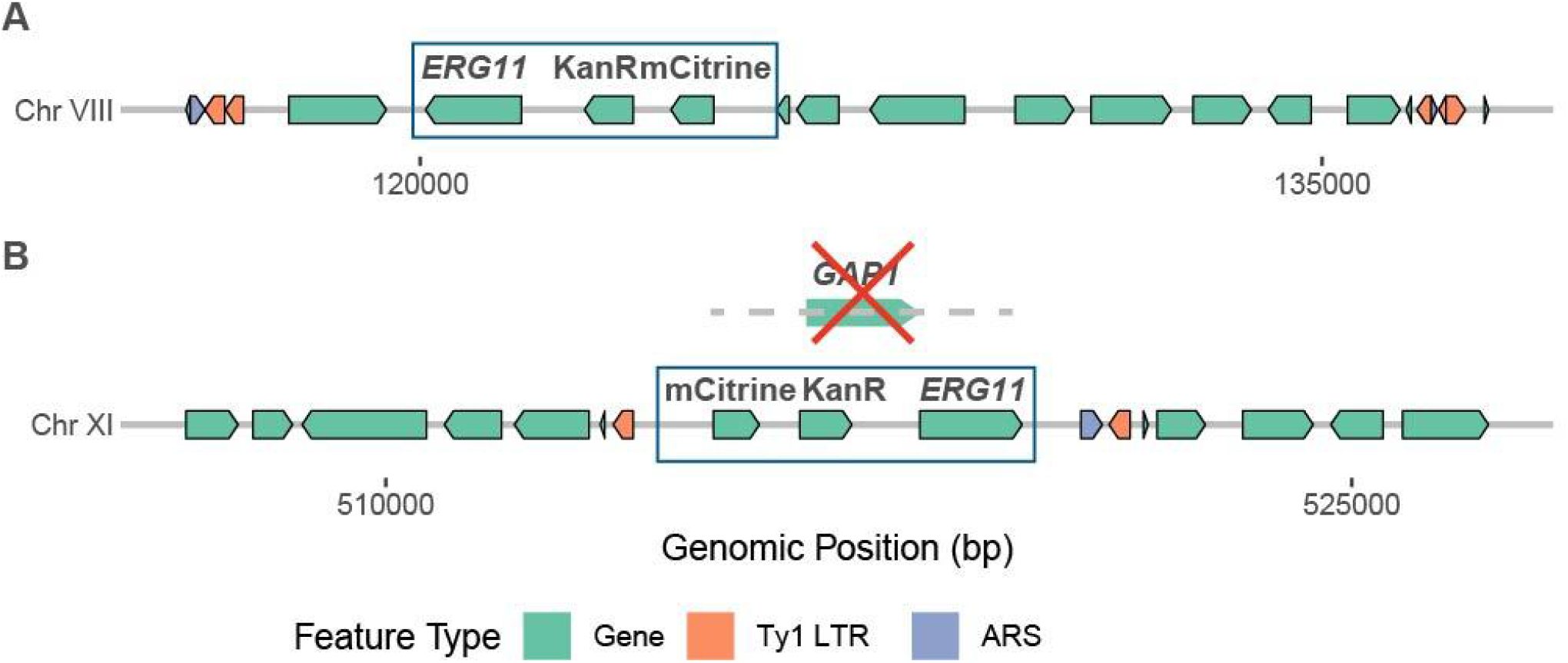
Genomic context of *ERG11* CNV reporter. **A**) The *ERG11* CNV reporter strain at the *ERG11* native locus on chromosome VIII and **B**) strain EKO_EaG at the *GAP1* locus on chromosome XI, showing surrounding features including genes, Ty1 LTR-retrotransposon elements, and autonomously replicating sequences. The mCitrine-KanR CNV reporter with the *ERG11* gene is highlighted.

**S2 Fig.**
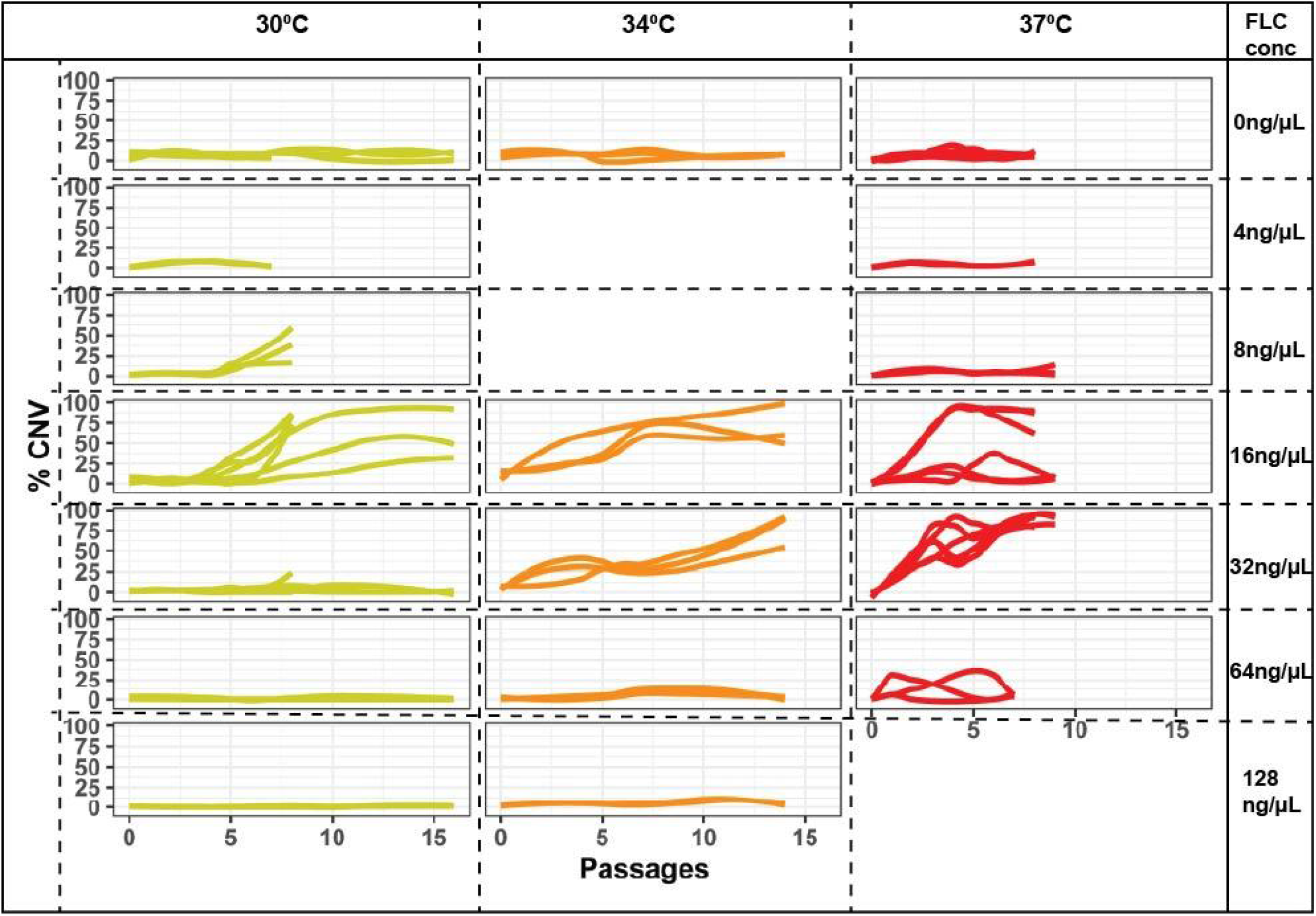
*ERG11* CNVs emerge in a narrow range of fluconazole concentrations in a temperature-dependent manner. Each line represents a single population of cells undergoing evolution by serial dilution in a given concentration of fluconazole (noted at the top of the graph). Line color denotes temperature at which the population was evolved. Data from all samples for all passages of serial dilution is shown.

**S3 Fig.**
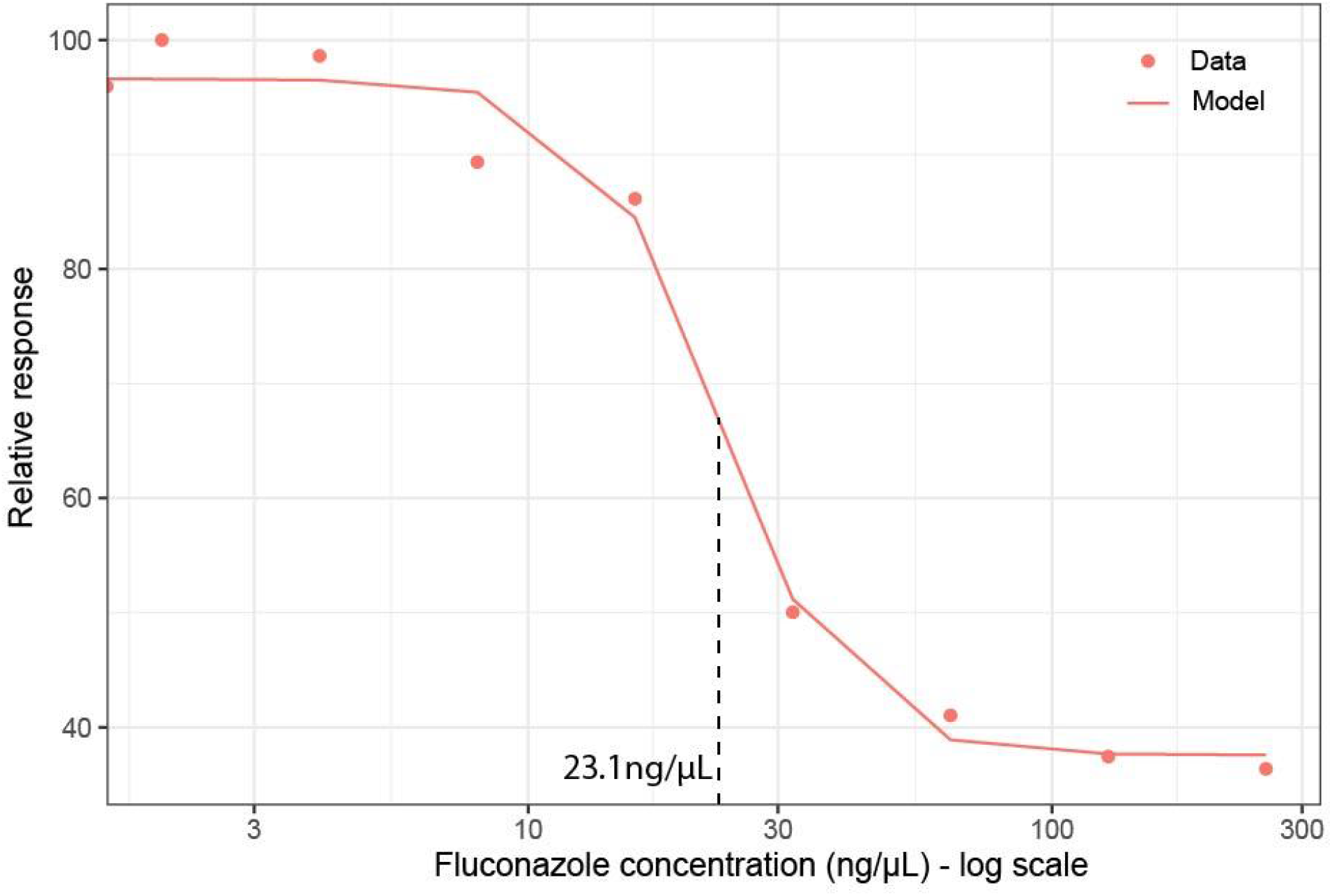
IC_50_ for fluconazole for *S. cerevisiae* in rich media is 23.1ng/μL. Dose-response curve for a total of 20 replicates naive *S. cerevisiae* strains (WT,1_ctrl, 2_ctrl, mCh_ctrl; see **S1 Table**) in varying concentrations of fluconazole, with relative growth rate (normalized to growth rate with no fluconazole) plotted on the y-axis.

**S4 Fig.**
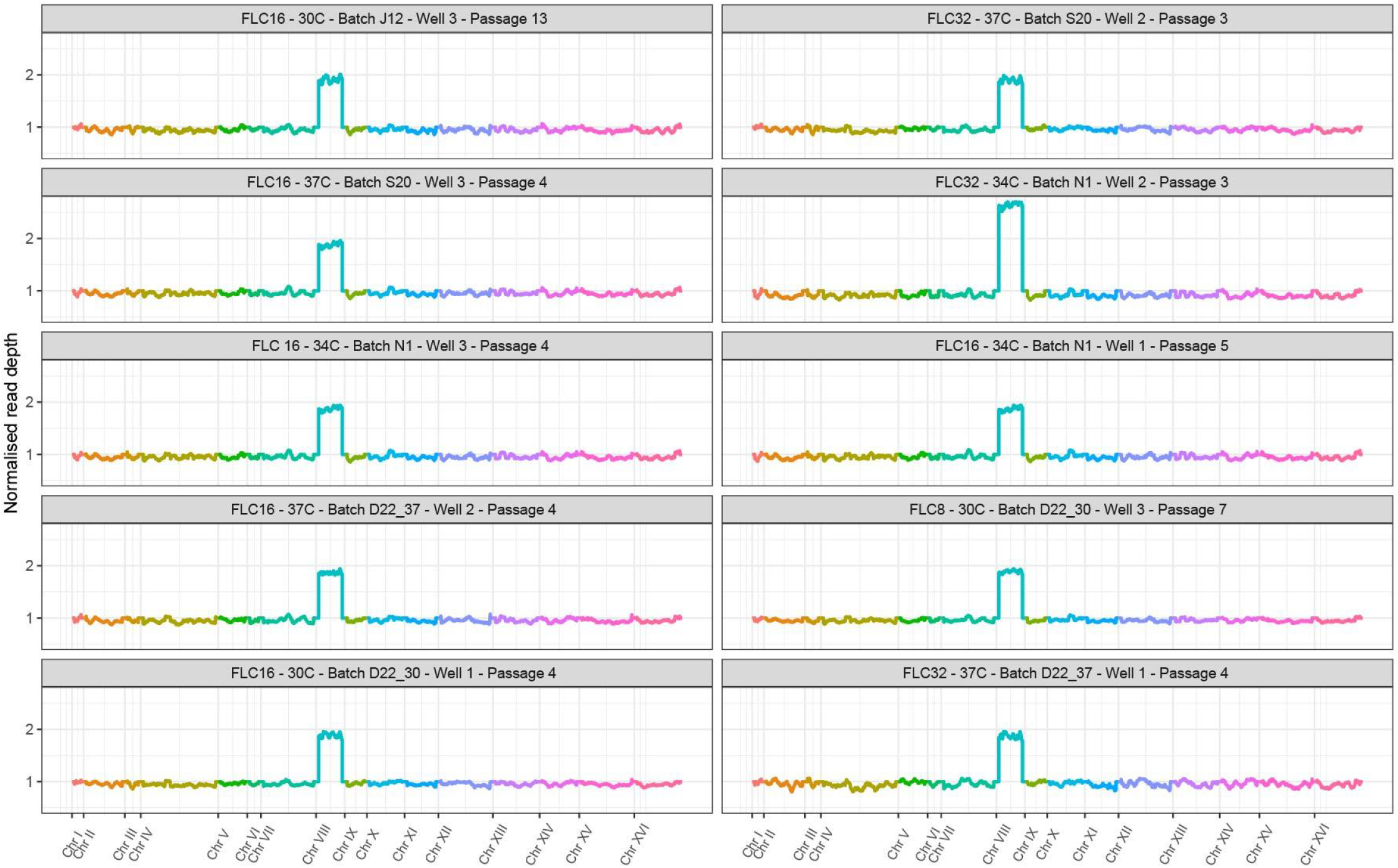
Sequenced *ERG11* CNV strains are aneuploidies. Genome-wide read depth of 10 representative *ERG11* CNV-containing strains, evolved in a variety of fluconazole concentrations and growth temperatures, as indicated in the title of each panel. All strains feature a full-chromosome duplication of chromosome VIII, while one (FLC32-34C) contains a triplication.

**S5 Fig.**
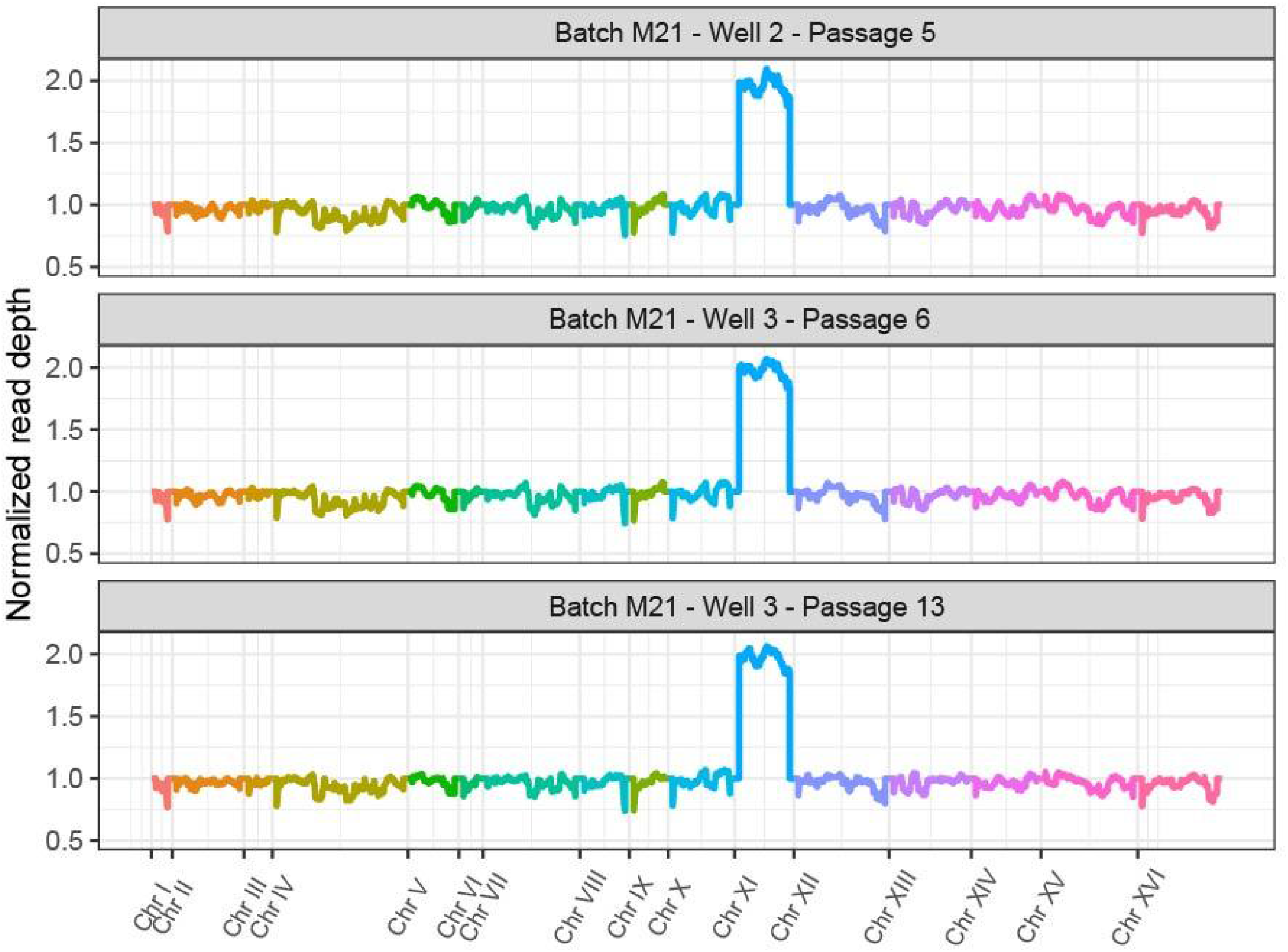
EaG_EKO CNV strains are chromosome XI aneuploidies. Genome-wide read depth 3 *ERG11* CNV-containing strains evolved at 16ng/μL fluconazole and 30℃.

**S6 Fig.**
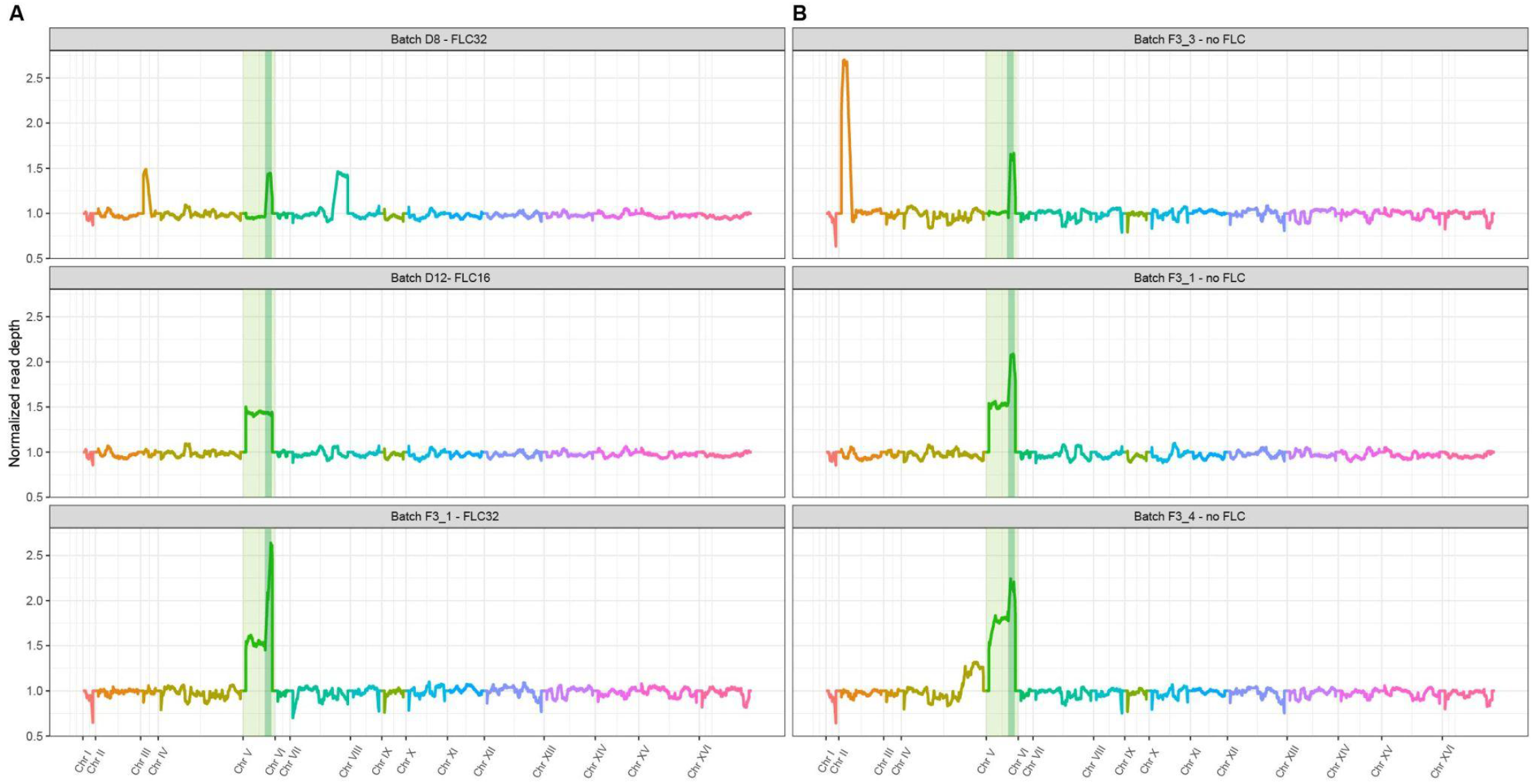
Sequenced FACu strains show CNV diversity. Genome-wide read depth of representative strains isolated after acute formaldehyde-copper sulphate selection after overnight growth in the (**A**) presence and (**B**) absence of fluconazole. chromosome V is highlighted in light green and the *SFA1-CUP1* locus is highlighted in dark green.

